# Dose Modeling for Mitochondrial Transplantation in Barrier-Limited Tissues

**DOI:** 10.1101/2025.07.08.663384

**Authors:** Melanie Walker, Giuseppe Orlando, Amish Asthana, Gabriel C. Oniscu, Michael R. Levitt

## Abstract

Mitochondrial transplantation is an emerging regenerative therapy aimed at preventing ischemia– reperfusion injury (IRI) by restoring oxidative metabolism in vulnerable tissues. While early clinical studies have demonstrated feasibility in pediatric myocardium and safety in adult stroke patients, dosing strategies remain undefined for barrier-limited organs where autoregulation, restricted perfusion, and anatomical constraints limit parenchymal uptake. We developed a physiologically grounded computational model to estimate route-specific input dose requirements for brain, spinal cord, and kidney. The model integrates organ-level parameters including tissue mass, regional perfusion or cerebrospinal flow, permeability–surface area product, and extraction efficiency. A benchmark dose of 2 × 10^6^ mitochondria per gram, derived from validated cardiac protocols, was applied across delivery scenarios. Sensitivity analysis explored the effects of extraction fraction and tissue mass. Route-adjusted input requirements diverged quantitatively from cardiac-derived estimates, with up to 2-fold increases in central nervous system (CNS) targets and approximately 20% higher dosing required for the kidney. This framework provides a quantitative foundation for translational dose planning in IRI and related conditions.

## INTRODUCTION

Mitochondrial transplantation has emerged as a candidate therapeutic strategy for IRI, which is characterized by bioenergetic failure, oxidative stress, and metabolic collapse ^1^. Preclinical studies have demonstrated that viable, respiration-competent mitochondria can be isolated and delivered to ischemic or metabolically impaired tissues, where they restore oxidative phosphorylation, augment ATP production, and mitigate cell death in models of myocardial infarction, acute kidney injury, spinal cord trauma, and neurodegenerative disease ^2–7^. Early-phase clinical studies in which autologous mitochondria were directly injected into the ischemic myocardium reduced IRI and improved outcomes in pediatric patients requiring extracorporeal membrane oxygenation ^8^. An additional ongoing trial is evaluating intra-arterial mitochondrial infusion in adults with acute ischemic stroke undergoing endovascular reperfusion therapy ^9^.

These studies have established procedural feasibility and safety, with preliminary evidence of biological activity and functional tissue rescue.

This study builds directly on the foundational cardiac work by McCully and colleagues ^5,10,11^, which demonstrated the efficacy and feasibility of mitochondrial transplantation in the pediatric myocardium. It extends those principles to adult non-cardiac tissues with distinct anatomical and procedural constraints. Myocardial delivery benefits from homogeneous perfusion, dense mitochondrial packing in cardiomyocytes, and direct or coronary injection access ^5^. In contrast, delivery to the CNS and kidney is constrained by anatomical barriers such as the blood–brain barrier (BBB), strict compartmental volume limits, and regional perfusion heterogeneity. These organs were selected as modeling targets based on their relevance in IRI and their shared property of intrinsic autoregulation of blood flow ^12^, which further restricts capillary-level uptake and delivery efficiency. By integrating anatomical constraints and route-specific delivery parameters, the framework supports quantitative dose planning for future clinical trials and may inform translation to other conditions involving regional mitochondrial dysfunction.

## METHODS

### Model Scope and Structure

This study developed a computational model to estimate mitochondrial dose requirements based on a functional restoration framework. Instead of replicating the full endogenous mitochondrial pool of a given tissue, the model defines a biologically sufficient dose aimed at restoring oxidative capacity to a therapeutically meaningful level. The benchmark dose of 2 × 10^6^ mitochondria per gram was derived from myocardial transplantation studies demonstrating efficacy in reducing infarct size and improving function ^5,13–16^. This benchmark was applied to the adult human brain, spinal cord, and kidney. Parenchymal cellularity estimates were taken from stereological and anatomical studies ^17–19^. Route-specific extraction fractions were assigned based on published capillary permeability–surface area and regional blood flow parameters ^20–22^. Calculated input doses were evaluated against injection volume constraints using experimentally validated suspension concentrations ^5,23^.

The model used a compartmental structure, with each simulation representing an independent anatomical target defined by a discrete vascular territory or a cerebrospinal fluid–accessible region. Four intra-arterial targets were modeled: the cortical and subcortical regions perfused by the right MCA via the M1 segment, a focal M2 branch territory, a single spinal cord segment supplied bilaterally by radiculomedullary arteries, and the renal cortex and medulla via the renal artery. Two intrathecal targets were included: the caudal spinal cord (lumbar puncture) and the basal cisterns (cisternal injection). Unless otherwise specified, each simulation reflects a single injection into a discrete anatomical compartment: unilateral intra-arterial delivery to the right MCA (M1 or M2), a single renal artery, or a single spinal segment; and focal intrathecal delivery via either cisternal or lumbar injection. These assumptions reflect typical procedural access points used in translational studies and provide a standardized basis for comparing route-specific dose requirements.

Simulations were based on adult human physiology, without pediatric scaling or animal extrapolation. Each began with a defined parenchymal dose, scaled by route-specific extraction efficiency to compute input dose and corresponding injection volume at the validated suspension limit. The model excluded recirculation and multicompartment kinetics, treating intra-arterial delivery as single-pass capillary uptake and intrathecal delivery as unidirectional CSF-to-parenchyma influx. Simulations were implemented in Python 3.10 using NumPy ^24^, SciPy ^25^, and pandas. Dose and injection volume calculations were performed using direct algebraic formulas. The complete set of parameter values and anatomical reference data is presented in Table S1.

### Benchmark Dose and Functional Restoration Threshold

The therapeutic dose target was defined as the number of viable mitochondria required per gram of parenchymal tissue to restore a biologically meaningful level of oxidative capacity. Rather than attempting to replicate the full endogenous mitochondrial population, the model adopted a benchmark threshold of 2 × 10^6^ mitochondria per gram, based on the upper end of the effective therapeutic range validated in myocardial mitochondrial transplantation studies ^13,26,27^. This benchmark was treated as generalizable to other highly perfused, metabolically active tissues, including the brain, spinal cord, and kidney.

For each modeled target, the total parenchymal dose was calculated as:

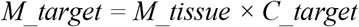

where *M_target* is the number of mitochondria required within the parenchyma, *M_tissue* is the modeled mass of the target tissue in grams, and *C_target* is the benchmark threshold (2 × 10^6^ mitochondria per gram). This approach assumes homogeneous distribution of therapeutic mitochondria within the viable parenchyma and does not account for regional heterogeneity in uptake or function.

### Route-Specific Extraction Modeling

To convert the benchmark tissue requirement into an administered input dose, each simulation applied a route-specific extraction fraction, *E*, defined as the proportion of administered mitochondria that successfully reach the parenchyma. For intra-arterial delivery, extraction was modeled using a physiologically validated capillary perfusion framework ^28–33^ in which uptake occurs across the endothelial barrier according to the Renkin-Crone ^34^ equation:

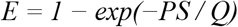

where *PS* is the permeability–surface area product in milliliters (mL) per minute, and *Q* is the regional blood flow in mL per minute. Although originally derived for passive solute diffusion, the Renkin–Crone equation is used here to approximate delivery-limited uptake of mitochondria.

In this context, the *PS* term reflects an effective clearance parameter, and the model assumes that uptake is limited by perfusion rather than by saturation of intracellular transport.

For MCA territory delivery, blood flow was set at 200 mL per minute ^20^. A *PS* value of 400 mL per minute represented an intact BBB, producing *E* ≈ 0.86. A value of 1000 mL per minute reflected BBB disruption, yielding *E* ≈ 0.99.

For the kidney, *Q* was set at 300 mL per minute and *PS* at 600 mL per minute ^35^, producing *E* ≈ 0.85. This value reflects uptake from the peritubular capillary network rather than glomerular filtration, which is not expected to occur due to the size of intact mitochondria ^36,37^. For spinal arterial delivery, extraction fractions were estimated using the same benchmark as for neural tissue, based on the physiological similarity between the blood–spinal cord barrier and the BBB, and due to a lack of validated spinal capillary PS/Q data in humans ^38^. Intrathecal delivery was modeled as a non-vascular route in which extraction reflects mitochondrial transit from cerebrospinal fluid to adjacent interstitial space. Extraction fractions were assigned based on published studies of cerebrospinal tracer distribution ^39–41^ and glymphatic transport ^42–44^. Lumbar intrathecal injection was assigned *E* = 0.30, consistent with limited tracer entry into caudal spinal tissue ^41–44^. Cisternal injection was assigned *E* = 0.50, based on more favorable uptake into brainstem and ventral forebrain regions ^45,46^.

In all simulations, the adjusted input dose was calculated as:

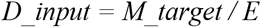

where *D_input* is the total number of mitochondria that must be administered to deliver the desired parenchymal dose. The assignment of extraction fractions in this model reflects both quantitative estimates from capillary perfusion equations and qualitative factors such as endothelial uptake and barrier integrity. Table S2 provides a conceptual summary of route-specific influences on extraction efficiency.

### Suspension Constraints and Volume Limits

To assess the practical feasibility of delivering the input dose, each simulation computed the injection volume required to deliver *D_input* at a maximum feasible suspension concentration of 2 × 10^8^ mitochondria per mL. This value is based on validated injectability thresholds for buffered mitochondrial preparations, beyond which aggregation, loss of viability, and delivery obstruction become likely ^27^. Required injection volume was computed using:

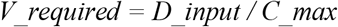

where *V_required* is the injection volume in mL and *C_max* is the suspension concentration ceiling of 2 × 10^8^ mitochondria per mL. Injection volumes were compared against route-specific feasibility thresholds. For intra-arterial delivery, a maximum volume of 10 mL was used, based on protocols for intracoronary ^23,47^ and intracerebral infusion ^48,49^. Fluid volume in the cerebral vasculature is a known contributor to intracranial pressure, which further limits the tolerable volume of infusion in neurovascular applications ^50^. For intrathecal injection, the upper limit was set at 2 mL, consistent with clinical safety parameters for lumbar and cisternal delivery ^51–53^.

Simulations exceeding these volumes were flagged as infeasible and interpreted as requiring further optimization or alternative delivery strategies.

### Sensitivity Analysis

To characterize model robustness and identify dose-limiting variables, a univariate sensitivity analysis was performed. Three parameters were varied independently: benchmark mitochondrial dose per gram, extraction fraction, and tissue mass. The benchmark dose was varied from 1 × 10^6^ to 3 × 10^6^ mitochondria per gram. Extraction fractions were varied from 0.50 to 0.99 for intra-arterial delivery and from 0.15 to 0.50 for intrathecal delivery, reflecting physiological ranges reported in tracer and contrast studies. Tissue mass was varied by ±15% around the reference values assigned for each target.

Simulations were implemented as parameter sweeps. For each value, the input dose and injection volume were recalculated, and feasibility was reassessed against fixed delivery thresholds. No probabilistic, multivariate, or interaction modeling was applied. All sensitivity outputs were stored in structured data frames for reproducibility.

## RESULTS

### Route-Adjusted Mitochondrial Dose Estimates Across Target Regions

Input dose requirements and associated injection volumes were calculated for six anatomically and clinically relevant targets using a benchmark therapeutic threshold of 2 × 10^6^ mitochondria per gram of viable parenchyma (Table 1).

**Table 1.** Modeled Mitochondrial Dose and Volume Requirements Across Routes and Targets at Varying Extraction Fractions.

| Route | Target Region | Target Mass (g) | Extraction Fraction (*E*) | Total Dose (mito) | Input Dose (mito) | Injection Volume* (mL) |
| --- | --- | --- | --- | --- | --- | --- |
| IA | MCA (M1) | 300 | 0.99 | $6.00 \times 10^8$ | $6.06 \times 10^8$ | 3.03 |
| IA | MCA (M1) | 300 | 0.80 | $6.00 \times 10^8$ | $7.50 \times 10^8$ | 3.75 |
| IA | MCA (M1) | 300 | 0.50 | $6.00 \times 10^8$ | $1.20 \times 10^9$ | 6.00 |
| IA | MCA (M2) | 50 | 0.99 | $1.00 \times 10^8$ | $1.01 \times 10^8$ | 0.51 |
| IA | MCA (M2) | 50 | 0.80 | $1.00 \times 10^8$ | $1.25 \times 10^8$ | 0.63 |
| IA | MCA (M2) | 50 | 0.50 | $1.00 \times 10^8$ | $2.00 \times 10^8$ | 1.00 |
| IA | Spinal Segment† | 1.5 | 0.99 | $3.00 \times 10^6$ | $3.03 \times 10^6$ | 0.02 |
| IA | Spinal Segment† | 1.5 | 0.80 | $3.00 \times 10^6$ | $3.75 \times 10^6$ | 0.02 |
| IA | Spinal Segment† | 1.5 | 0.50 | $3.00 \times 10^6$ | $6.00 \times 10^6$ | 0.03 |
| IA | Kidney | 300 | 0.85 | $6.00 \times 10^8$ | $7.06 \times 10^8$ | 3.53 |
| IT | Lumbar | 1.5 | 0.30 | $3.00 \times 10^6$ | $1.00 \times 10^7$ | 0.05 |
| IT | Cisternal | 1.5 | 0.50 | $3.00 \times 10^6$ | $6.00 \times 10^6$ | 0.03 |
| **Key:** IA, intra-arterial; IT, intrathecal, MCA, middle cerebral artery; mito, mitochondria.<br>* All injection volumes assume a suspension concentration of $2.00 \times 10^8$ mitochondria/mL.<br>† For the spinal segment via intra-arterial (IA) route, injection volumes are reported per side. Bilateral delivery would require doubling the stated volume. |  |  |  |  |  |  |

For M1 cortical delivery, input dose requirements ranged from 6.06 × 10^8^ to 1.20 × 10^9^ mitochondria, corresponding to injection volumes between 3.03 and 6.00 mL. In the smaller M2 territory, volumes ranged from 0.51 to 1.00 mL across the same extraction range. For spinal and renal targets, modeled input doses remained within procedural feasibility thresholds. The spinal segment simulation assumed bilateral infusion and a combined parenchymal mass of 1.5 grams.

For intrathecal delivery, required input doses were lower but constrained by limited extraction efficiency, resulting in injection volumes of 0.05 mL (lumbar) and 0.03 mL (cisternal).

### Sensitivity to Benchmark Dose and Target Mass

Sensitivity simulations evaluated the effect of varying the benchmark mitochondrial dose from 1 × 10^6^ to 3 × 10^6^ mitochondria per gram and adjusting tissue mass ±15%. Extraction fractions were fixed at route-specific values: *E* = 0.99 for cortical delivery, *E* = 0.85 for kidney, *E* = 0.80 for spinal intra-arterial, and *E* = 0.30 for intrathecal administration.

Both benchmark dose and tissue mass scaled linearly with input dose and injection volume. For M1 cortex, halving the benchmark to 1 × 10^6^ reduced volume from 3.75 to 1.88 mL, while increasing it to 3 × 10^6^ raised volume to 5.63 mL. A ±15% change in tissue mass produced proportional changes in all target regions. Modeled conditions remained within procedural volume limits across the parameter space. Results are provided in Table S3.

### Comparison to Cardiac Dosing

To assess the value of route-aware modeling, dose requirements were compared to those generated by direct extrapolation of the validated cardiac protocol, in which 2 × 10^6^ mitochondria per gram were delivered without adjustment for delivery route or extraction efficiency.

Under high-efficiency conditions (*E* = 0.99), route-adjusted doses closely matched cardiac-derived values. At lower extraction fractions (*E* = 0.80 or 0.50), input doses increased by up to 100%, with corresponding increases in injection volume. Differences were most pronounced in CNS and intrathecal targets. Table 2 summarizes the magnitude of divergence between cardiac-derived and route-adjusted estimates for all modeled regions.

**Table 2.**
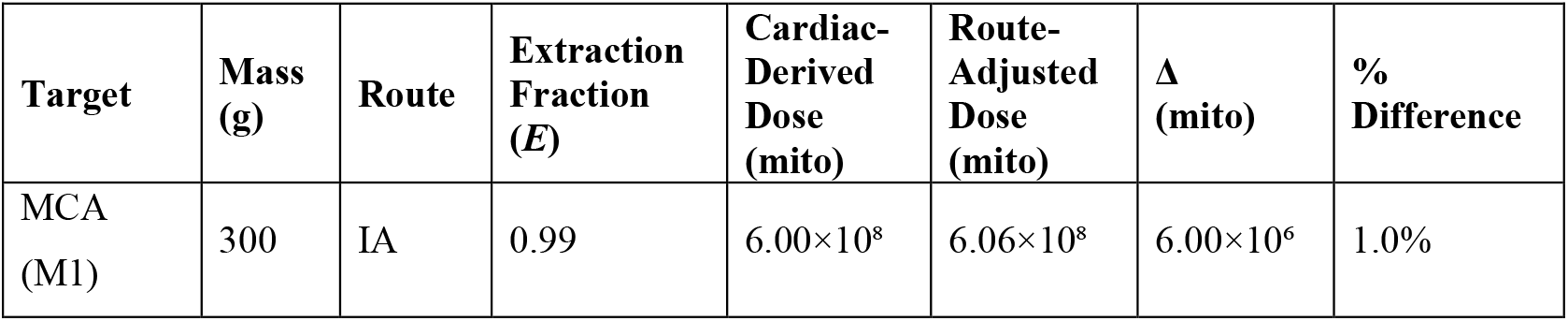
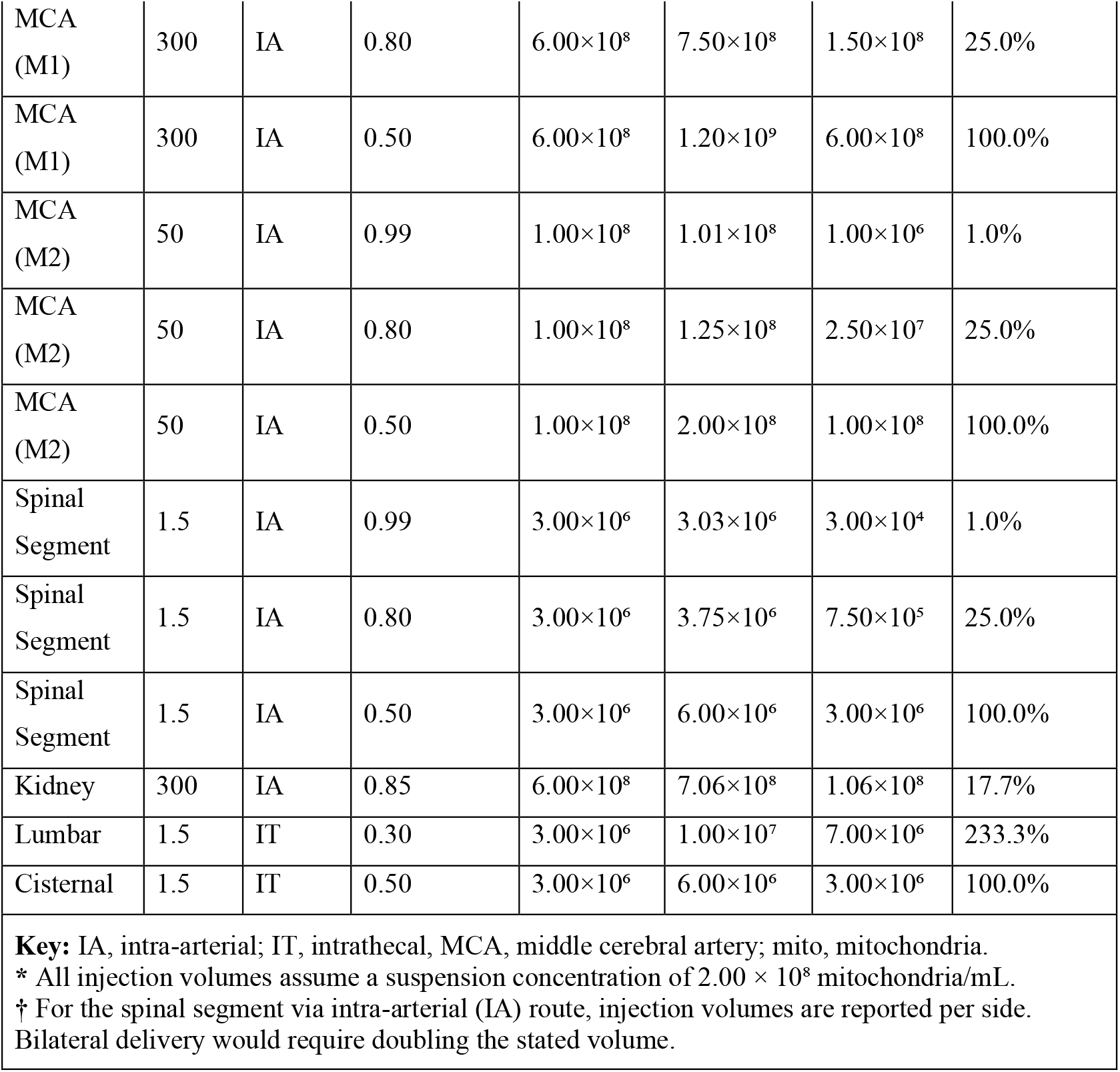
Comparison of Route-Adjusted vs. Cardiac-Derived Mitochondrial Dose Estimates.

## DISCUSSION

Building on validated cardiac transplantation models ^10,13,26,27,47^, this study presents a physiologically grounded framework for mitochondrial dose planning in tissues with more complex anatomical barriers, regional delivery constraints, or heterogeneous perfusion dynamics. Simulation results demonstrate that extraction efficiency significantly alters the relationship between administered and effective dose, depending on the delivery route and anatomical barriers. In the myocardium, the benchmark dose of 2 × 10^6^ mitochondria per gram is deliverable either by direct injection or coronary perfusion, aided by dense mitochondrial content and high capillary surface area in cardiomyocytes ^16,54^. These conditions enable efficient uptake with minimal loss, supporting the benchmark dose used for other tissues ^5,16^. In contrast, CNS tissues are characterized by heterogeneous perfusion, restricted extracellular space, and barrier systems such as the BBB and arachnoid membrane ^43,51,55–57^, all of which constrain mitochondrial entry and volume tolerability. Failing to account for these factors may result in suboptimal dosing when extrapolating from existing protocols, potentially causing under delivery or requiring excessive injection volumes.

The Renkin–Crone equation, originally developed to describe passive solute diffusion using the indicator-diffusion method ^34^, has since been applied widely in tissue perfusion modeling across organ systems including brain, kidney, and liver ^20,21,32,58^. In contemporary imaging studies ^59^, extraction-corrected *PS* models are used to estimate delivery-limited uptake of contrast agents without invoking equilibrium diffusion. We adopt this formulation in the present study to represent mitochondrial delivery-limited uptake, rather than to describe mechanistic permeability or intracellular transport. In this context, the *PS* parameter reflects an effective clearance term governed by capillary surface area, regional blood flow, and barrier integrity, without explicitly modeling internalization kinetics.

Unlike traditional therapeutics, mitochondria are metabolically active, double-membraned organelles that generate ATP, regulate redox balance, and interact dynamically with host cell networks ^60^. Their size (approximately 0.5 to 1 micrometer) and functional complexity distinguish them from inert drugs or biologic macromolecules ^60^. Although intact extracellular mitochondria have been detected in the human circulation ^61^, their biodistribution does not follow classical pharmacokinetic rules ^62,63^. They are not filtered by the kidney ^64,65^, do not diffuse across vascular beds passively ^66,67^, and lack a definable or constant systemic half-life ^68–70^. Therapeutic efficacy depends on targeted delivery, successful cellular uptake, and retention of metabolic function within recipient cells. As a result, dose-response relationships are anatomically localized, route dependent, and determined primarily by physical access rather than systemic exposure. Conventional pharmacokinetic parameters such as volume of distribution or area under the curve are not applicable to mitochondrial delivery.

Intra-arterial delivery to CNS territories with an intact BBB results in low extraction efficiency and necessitates higher administered doses to meet parenchymal thresholds. Barrier disruption in stroke or trauma improves access, but endothelial sequestration may reduce net delivery.

Although not explicitly modeled, this could explain observed variability in uptake. Spinal cord capillary properties are less well characterized ^38^, but focal intra-arterial targeting appears viable within volume constraints. Intrathecal injection bypasses vascular barriers but is limited by slow glymphatic influx and regional flow dynamics ^42–44^, resulting in reduced extraction fractions and correspondingly higher dose requirements for a given effect.

Comparison with cardiac-derived extrapolations supports the importance of correcting for delivery routes. When extraction is near-complete (*E* = 0.99), the benchmark model and cardiac estimates converge. At lower extraction values (*E* = 0.50 or 0.80), the divergence can exceed twofold, with injection volumes approaching or exceeding clinical limits. These findings emphasize the need for route-specific dose planning in tissues with delivery barriers and highlight the limitations of empirical extrapolation.

This model has several limitations related to biological complexity and delivery assumptions. It was designed to estimate the minimum number of viable mitochondria required in the injectate to achieve functional parenchymal delivery, under anatomically and procedurally constrained conditions. The model does not account for downstream biological processes that may influence efficacy or biodistribution, including intracellular trafficking, fusion with endogenous mitochondrial networks, or organelle degradation. Losses related to catheter retention, adhesion to delivery systems, or aggregation in suspension were also not included. Intravenous (IV) delivery was not modeled, as prior animal studies have shown that fewer than one percent of systemically administered mitochondria reach the brain, spinal cord, or kidney, with the majority sequestered in the lungs and liver ^71–73^. Although other classes of mitochondrial-targeted therapies may be systemically viable, these constraints make IV administration unsuitable for achieving therapeutic concentrations of exogenously delivered mitochondria in barrier-limited tissues

## CONCLUSION

Ischemic vascular disease is the leading contributor to global disability and mortality ^74^, yet no therapies exist to directly treat IRI. Early trials of mitochondrial transplantation have demonstrated biological feasibility and initial clinical promise. However, no standardized or physiology-based approach exists for predicting dosing in tissues with complex anatomical features such as autoregulation, restricted perfusion, or compartment-specific access. This study provides a reproducible, anatomy-based framework for estimating mitochondrial dose in such contexts, incorporating route-specific delivery parameters. These results support tailored trial design for IRI and offer a foundation for future translational applications.

## ACKNOWLEDGEMENTS

This research was conducted without public, private, or commercial funding support.

## AUTHOR CONTRIBUTIONS

MW was responsible for study design, computational analysis, and manuscript drafting. All authors provided technical input, participated in data review, and contributed to manuscript revision. All authors approved the final version of the manuscript for submission.

## DATA AVAILABILITY

All data generated or analyzed during this study are included in this published article and its supplementary information files.

## CODE AVAILABILITY

The underlying Python code is available in the supplementary information files.

