## Supplemental File for "Dose Modeling for Mitochondrial Transplantation in Barrier-Limited Tissues"

**Supplementary Information**

**Supplemental Tables S1-S3**

Table S1. Modeled Delivery Parameters and Benchmark Dose Assumptions Across Routes and Anatomical Targets

| Delivery Route | Target Region | Target Mass (g) | Benchmark Dose (# mito) | Extraction Fraction (*E*) | Max Volume (mL) | Notes |
|---|---|---|---|---|---|---|
| IA | MCA (M1) | 300 | $6.00 \times 10^8$ | 0.50–0.99 (open BBB) | 10 | Modeled with BBB disruption data[20] |
| IA | MCA (M2) | 50 | $1.00 \times 10^8$ | 0.50–0.99 (open BBB) | 5 | Approximates branch territory as M1[20,21] |
| IA | Spinal Segment | 1.5 | $2 \times 10^6$ mito/g | 0.50–0.99 (benchmark) | 10 (5 per side) | Bilateral radiculomedullary arteries[75,76] |
| IA | Kidney | 300 | $6.00 \times 10^8$ | 0.85 | 10 | Full renal artery perfusion[22,35] |
| IT | Lumbar | 50 | $1.00 \times 10^8$ | 0.15–0.30 | 2 | Based on tracer and glymphatic uptake in caudal spinal cord[43] |
| IT | Cisternal | 50 | $1.00 \times 10^8$ | 0.30–0.50 | 2 | Enhanced uptake into brainstem and forebrain[45,46] |
| Key: IA, Intra-arterial; IT, intrathecal; mito, mitochondria; BBB, blood-brain barrier | | | | | | |

**Table S2. Conceptual Influence of Barrier Integrity and Endothelial Capture on Extraction Fraction**

| Delivery Route | BBB Status / Barrier Type | Endothelial Capture | Approximate *E* | Key Limiting Factor(s) |
|---|---|---|---|---|
| IA | Intact (normal) | High | 0.01 – 0.10 | Tight junctions, poor permeability, high endothelial uptake |
| IA | Disrupted (stroke) | Moderate | 0.80 – 0.99 | High *PS*, open junctions, increased parenchymal access |
| IA | Disrupted BBB with high endothelial capture | High | 0.60 – 0.80 | Endothelial uptake reduces effective despite high permeability |
| IA | Renal glomerular and tubulointerstitial beds | Moderate | ~0.85 | Capillary fenestration, high perfusion, moderate non-parenchymal loss |
| IT | Bypassed (nonvascular CSF–parenchyma) | N/A | 0.15 – 0.30 | Limited glymphatic access, slow interstitial penetration |
| IT | Bypassed (CSF–parenchyma) | N/A | 0.30 – 0.50 | Favorable access to brainstem and ventral forebrain via basal cisterns |
| **Key:** BBB, blood-brain barrier; CSF, cerebrospinal fluid; IA, Intra-arterial; IT, intrathecal | | | | |

**Table S3. Sensitivity of Dose and Injection Volume to Benchmark Threshold and Target Mass Under Fixed Extraction Efficiency**

| Route | Target Region | Target Mass (g) | Extraction Fraction (*E*) | Benchmark Dose (mito/g) | Total Dose (mito number) | Input Dose (mito number) | Injection Volume (mL) |
|---|---|---|---|---|---|---|---|
| IA | MCA (M1) | 300 | 0.99 | $1.00\times10^6$ | $3.00\times10^8$ | $3.03\times10^8$ | 1.52 |
| IA | MCA (M1) | 300 | 0.99 | $2.00\times10^6$ | $6.00\times10^8$ | $6.06\times10^8$ | 3.03 |
| IA | MCA (M1) | 300 | 0.99 | $3.00\times10^6$ | $9.00\times10^8$ | $9.09\times10^8$ | 4.55 |
| IA | MCA (M1) | 255 | 0.99 | $2.00\times10^6$ | $5.10\times10^8$ | $5.15\times10^8$ | 2.58 |
| IA | MCA (M1) | 345 | 0.99 | $2.00\times10^6$ | $6.90\times10^8$ | $6.97\times10^8$ | 3.49 |
| IA | Kidney | 300 | 0.85 | $1.00\times10^6$ | $3.00\times10^8$ | $3.53\times10^8$ | 1.77 |
| IA | Kidney | 300 | 0.85 | $2.00\times10^6$ | $6.00\times10^8$ | $7.06\times10^8$ | 3.53 |
| IA | Kidney | 300 | 0.85 | $3.00\times10^6$ | $9.00\times10^8$ | $1.06\times10^9$ | 5.30 |
| IA | Kidney | 255 | 0.85 | $2.00\times10^6$ | $5.10\times10^8$ | $6.00\times10^8$ | 3.00 |
| IA | Kidney | 345 | 0.85 | $2.00\times10^6$ | $6.90\times10^8$ | $8.12\times10^8$ | 4.06 |
| IA | Spinal Segment | 1.5 | 0.80 | $2.00\times10^6$ | $3.00\times10^6$ | $3.75\times10^6$ | 0.02 |
| IT | Cisternal | 1.5 | 0.15 | $2.00\times10^6$ | $3.00\times10^6$ | $2.00\times10^7$ | 0.10 |
| IT | Cisternal | 1.5 | 0.50 | $2.00\times10^6$ | $3.00\times10^6$ | $6.00\times10^6$ | 0.03 |
| IT | Cisternal | 1.3 | 0.30 | $2.00\times10^6$ | $2.60\times10^6$ | $8.67\times10^6$ | 0.04 |
| IT | Cisternal | 1.7 | 0.30 | $2.00\times10^6$ | $3.40\times10^6$ | $1.13\times10^7$ | 0.06 |
| IT | Cisternal | 1.5 | 0.30 | $1.00\times10^6$ | $1.50\times10^6$ | $5.00\times10^6$ | 0.03 |
| IT | Lumbar | 1.5 | 0.30 | $3.00\times10^6$ | $4.50\times10^6$ | $1.50\times10^7$ | 0.08 |
| IT | Lumbar | 1.3 | 0.30 | $2.00\times10^6$ | $2.60\times10^6$ | $8.67\times10^6$ | 0.05 |
| IT | Lumbar | 1.7 | 0.30 | $2.00\times10^6$ | $3.40\times10^6$ | $1.13\times10^7$ | 0.06 |
| IT | Lumbar | 1.5 | 0.30 | $2.00\times10^6$ | $3.00\times10^6$ | $1.00\times10^7$ | 0.05 |

**Key:** IA, Intra-arterial; IT, intrathecal; mito, mitochondria
* All injection volumes assume a suspension concentration of $2.00 \times 10^8$ mitochondria/mL.
† For the spinal segment via intra-arterial (IA) route, injection volumes are reported per side. Bilateral delivery would require doubling the stated volume.

**Python Script**

```
### Python Script: mito_dose_model.py

import numpy as np
import pandas as pd

# Constants
C_MAX = 2e8  # Maximum suspension concentration (mitochondria/mL)

# Define a function to calculate total dose, input dose, and injection
volume
def calculate_dose_and_volume(mass_g, benchmark_dose, E):
    total_dose = mass_g * benchmark_dose
    input_dose = total_dose / E
    volume_ml = input_dose / C_MAX
    return total_dose, input_dose, volume_ml

# Define the scenarios for sensitivity analysis
scenarios = [
    # (Route, Target, Mass (g), E, Benchmark Dose (mito/g))
    ('IA', 'MCA (M1)', 300, 0.99, 1e6),
    ('IA', 'MCA (M1)', 300, 0.99, 2e6),
    ('IA', 'MCA (M1)', 300, 0.99, 3e6),
    ('IA', 'MCA (M1)', 255, 0.99, 2e6),
    ('IA', 'MCA (M1)', 345, 0.99, 2e6),
    ('IA', 'Kidney', 300, 0.85, 1e6),
    ('IA', 'Kidney', 300, 0.85, 2e6),
    ('IA', 'Kidney', 300, 0.85, 3e6),
    ('IA', 'Kidney', 255, 0.85, 2e6),
    ('IA', 'Kidney', 345, 0.85, 2e6),
    ('IA', 'Spinal Segment', 1.5, 0.8, 2e6),
    ('IT', 'Cisternal', 1.5, 0.15, 2e6),
    ('IT', 'Cisternal', 1.5, 0.50, 2e6),
    ('IT', 'Cisternal', 1.3, 0.30, 2e6),
    ('IT', 'Cisternal', 1.7, 0.30, 2e6),
    ('IT', 'Cisternal', 1.5, 0.30, 1e6),
    ('IT', 'Lumbar', 1.5, 0.30, 3e6),
    ('IT', 'Lumbar', 1.3, 0.30, 2e6),
    ('IT', 'Lumbar', 1.7, 0.30, 2e6),
    ('IT', 'Lumbar', 1.5, 0.30, 2e6),
]

# Perform calculations and store results
```

```
results = []
for route, target, mass_g, E, benchmark_dose in scenarios:
    total_dose, input_dose, volume_ml =
calculate_dose_and_volume(mass_g, benchmark_dose, E)
    results.append({
        'Route': route,
        'Target Region': target,
        'Target Mass (g)': mass_g,
        'Extraction Fraction (E)': E,
        'Benchmark Dose (mito/g)': benchmark_dose,
        'Total Dose (mito)': total_dose,
        'Input Dose (mito)': input_dose,
        'Injection Volume (mL)': volume_ml
    })

# Create DataFrame and save
df = pd.DataFrame(results)
df.to_csv("/mnt/data/mito_dose_sensitivity_results.tsv", sep='\t',
index=False)
```

```
### README

# README: Mitochondrial Dose Sensitivity Simulation

## Purpose
This script models the input dose and injection volume required to
deliver viable mitochondria to specific human tissues. It accounts for
anatomical target mass, benchmark mitochondrial dose, and extraction
efficiency for intra-arterial (IA) and intrathecal (IT) routes.

## How It Works
Each simulation scenario specifies:
- Route (IA or IT)
- Target region
- Target tissue mass (grams)
- Benchmark mitochondrial dose per gram of tissue
- Extraction fraction (E)

Using these inputs, the script calculates:
- Total dose = Target Mass × Benchmark Dose
- Input dose = Total Dose / E
- Injection volume = Input Dose / Maximum Suspension Concentration

## Outputs
```

The script outputs a tab-separated values (.tsv) file named:
- `mito_dose_sensitivity_results.tsv`

This file includes all modeled scenarios described in the associated manuscript and supplemental Table S1.

### Requirements
- Python 3.10+
- NumPy
- Pandas

### Execution
To run:
```bash
python mito_dose_model.py
```

Make sure required packages are installed:
```bash
pip install numpy pandas
```
